# Epidemiology, Microbiology and Therapeutic Consequences of Chronic Osteomyelitis in Northern China: A Retrospective Analysis of 255 Patients

**DOI:** 10.1101/316836

**Authors:** Xianzhi Ma, Shengshou Han, Jun Ma, Xiaotao Chen, Wenbin Bai, Wenqi Yan, Kai Wang

## Abstract

The study aimed to explore the epidemiology and clinical characteristics of chronic osteomyelitis observed in a northern China hospital. Clinical data of 255 patients with chronic osteomyelitis from January 2007 to January 2014 were collected and analyzed, including general information, disease data, treatment and follow-up data. Chronic osteomyelitis is more common in males and in the age group from 41-50 years of age. Common infection sites are the femur, tibiofibular, and hip joint. More g+ than g- bacterial infections were observed, with *S. aureus* the most commonly observed pathogenic organism. The positive detection rate from debridement bacterial culture is 75.6%. The detection rate when five samples are sent for bacterial culture is 90.6%, with pathogenic bacteria identified in 82.8% of cases. The two-stage debridement method (87.0%) has higher first curative rate than the one-stage debridement method (71.2%). To improve detection rate using bacterial culture, at least five samples are recommended. Treatment of chronic osteomyelitis with two-stage debridement, plus antibiotic-loaded polymethylmethacrylate (PMMA) beads provided good clinical results in this study and is therefore recommended.

---

Chronic osteomyelitis is a common clinical disease and a challenging disorder, characterized by its long disease course, difficult early diagnosis and high disability rate. The clinical characteristics of chronic osteomyelitis are varied, and may be affected by geography, time, and pathogenetic differences. Geographically, developing countries have a higher incidence of the disease than developed ones, likely caused by differences in economic foundation, lifestyle and healthcare level [1-2]. Over time, a shift has occurred from predominately hematogenous osteomyelitis several decades ago, to a predominance of chronic osteomyelitis that results from trauma, implant infection, and diabetes [3-6].

In recent years, the number of patients with open wounds and multiple fractures from road and industrial accidents has sharply increased in China, as the Chinese economy undergoes rapid development. Multiple injuries are difficult to treat and frequently lead to infection; thus, treatment exerts great pressures on patients, both economically and psychologically. It also poses great challenges for orthopedists [7]. Wang [8] and Jiang et al. [9] provide recent data on the epidemiology of chronic osteomyelitis in southwest and southern China, which can be used by local government policy-makers and by clinicians. However, China is a country of great diversity in its population, climate and culture. Currently, data and relevant research are lacking for northern China on the epidemiology of chronic osteomyelitis. To address this, we conduct an epidemiological statistical analysis on 255 patients at a prominent hospital and explore the clinical characteristics of chronic osteomyelitis in northern China.

## Methods

A retrospective analysis was completed on 255 cases of patients with chronic osteomyelitis seen in the trauma orthopedic department at our hospital from January 2007 to January 2014. Patient inclusion criteria were:

(1) osteomyelitis diagnosis;
(2) local swelling and bone pain on examination;
(3) draining fistula present;
(4) imaging procedures completed;
(5) microbiological and histopathological examinations completed;
(1) biochemical examinations completed.

Exclusion criteria were:

(1) acute osteomyelitis (less than 2 weeks);
(2) no surgical treatment done;
(3) osteomyelitis site in the spine, pelvis or skull;
(4) complete data and examinations not available;
(5) patient had diabetic foot infection and was not treated in our department.

All data were obtained from the case management system in the medical record room of the hospital prior to this research.

The following data were collected:

1. Demographic information: age and gender;
2. Disease data: infection site, Cierny-Mader classification, laboratory examination results (including erythrocyte sedimentation rate (ESR), C-Reactive protein (CRP) and white blood cell (WBC)), and bacterial culture results (respectively from debridement and sinus tract);
3. Treatment method: two-stage debridement and one-stage debridement;
4. Follow-up data were obtained by searching medical records and by telephone. All patients were successfully followed-up 9-93 months (46.2 month in average) after the operation.

### Bacterial Culture and Identifying Pathogenic Bacteria

Different samples were collected during debridement and in sinus tract and sent for bacterial culture. The results are essential to identify pathogenic bacterial.

If the same bacteria type was detected in at least two different samples, it was identified as a pathogenic bacterium. Based on this principle, we examined the number of samples that tested positive with the number of samples analyzed, and the differences between them in identifying the pathogenic bacteria.

Bacterial cultures from the sinus tract samples can be an additional aid when examining chronic osteomyelitis and its result can serve as a reference for diagnosis and treatment. Samples from 58 patients with sinus tracts were sent for bacterial culture.

The coincidence rate between sinus tract positive result and identified pathogenic bacterial were examined, to assess its ability in identifying pathogenic bacteria.

### Treatment Methods

Two treatments were used in the study.

#### Two-stage Debridement

Two-stage debridement + Antibiotic-loaded

polymethylmethacrylate (**PMMA**) spacers ± External fixation

Step 1: Debridement and implantation of antibiotic-loaded PMMA. At the first stage of debridement, the antibiotics vancomycin and imipenem are used locally.
Step 2: During the one-week clinical observation, X-rays and CT scans are repeated, along with laboratory tests of CRP and ESR levels and reexamined while awaiting pathology results from the debridement bacterial culture. Patients received intravenous antibiotics during this time.
Step 3: Debridement was performed again after one week. The original antibiotic-impregnated cement beads were removed. Based on the pathogenic bacteria detected and its sensitivity test, appropriate antibiotic PMMA beads were implanted.

#### One-stage debridement

Debridement + Antibiotic-loaded PMMA spacers ± external fixation This treatment included only step 1 of the two-stage treatment above.

In the current study, we compared and analyzed their clinical effects and prognosis of each treatment.

##### Follow-up

In a follow-up session, clinical cure was assessed based on the following information: whether clinical symptoms of infection had disappeared, including local redness, swelling, heat, pain, abscess, and sinus tract; whether laboratory examination results no longer had indicators of infection, including WBC count, ESR, and CRP; and, whether progressive periosteum thickening or osteolysis was no longer present on a recent X-ray image.

##### Ethics approval and informed consent

The current study was approved by the ethical medical committee of the hospital. All participants gave informed written consent.

### Statistics Analysis

Statistical analysis was performed with SPSS software (Version 21.0; SPSS Inc., Chicago, IL). Frequency data were compared by Pearson’s chi-square test. For normally distributed data, independent groups were compared by a Student t-test or one-way analysis of variance (ANOVA). Where data were not normally distributed, the Mann–Whitney U test or Kruskal–Wallis H test was adopted. Results were considered significant at p < 0.05.

## Results

### Gender Ratio and Age at First Diagnosis

The present study included 202 males (79.2%) and 53 females (20.3%), an approximate gender ratio of 4:1.

The median age at first diagnosis was 45.5 years. Approximately 80% of affected patients were between 21 and 60 years of age (206 cases). The top three age groups represented were 41 to 50 years (29%), 21 to 30 years (20.8%), and 31 to 40 years (16.5%).

### Infection Site

All cases included in the current study were single-site infections. Among them, there were 77 cases of femur infections (30.2%), 66 tibiofibular infections (25.9%), 39 hip joint infections (15.3%), 16 ankle joint infections (6.3%), 14 humerus infections (5.5%), 14 ulna and radius infections (5.5%), 10 patella infections (3.9%), eight calcaneus infections (3.1%), five elbow infections (2.0%), four pelvis infections (1.6%), and two astragalus infections (0.8%).

### Classifications

All 255 cases met the Cierny-Mader classification criteria for chronic osteomyelitis (Table 1). Type IIA (23.1%), type IIIA (20.4%), and type IVA (18.0%), are the most frequent classifications in the study.

### Laboratory Test

The white blood cell count (WBC), ESR, and CRP of 255 patients before operation are presented in Table 2 below.

### Comprehensive Statistics

All data on WBC, ESR, and CRP were analyzed. The 255 patients had a total of 323 hospital admissions and WBC, ESR, and CRP were tested 304 times.

ESR and CRP are important indicators of chronic osteomyelitis in laboratory examination. If both ESR and CRP are abnormal, it is highly indicative of infection. If only CRP is increased, this may indicate infection. According to the results above, ESR and CRP were abnormal in 71.1% patients. Only 17.4% patients had an abnormal CRP result (Table 3).

### Pathogenic Microorganism

Bacterial culture results were classified into g+ bacteria, g- bacteria, or fungus. 245 samples were sent to bacterial culture. According to the result, g+ bacteria (160 times, 65.3%) were overall more common than g- bacteria (83 times, 33.9%). Fungus only appeared two times (0.8) in the result.

In total, 41 types of bacteria were detected. The 13 most frequently detected bacteria are given in Table 4.

Another 28 bacteria were detected three times or less, including *Klebsiella oxytoca* (3 times or 1.22%), *Citrobacter braakii* (3 times or 1.22%), *Staphylococcus sciuri* (2 times or 0.82%), *S. kloosii* (2 times or 0.82%), and *Proteus mirabilis* (2 times or 0.82%). The following bacteria were detected only once (0.41%): *A. calcoaceticus-A. baumannii* complex, *Achromobacter* spp., *Bacillus* sp., *Burkholderia cepacia*, *Candida parapsilosis*, *Corynebacterium minutissimum*, *Corynebacterium* sp., *Corynebacterium striatum*, *Dermatococcus* sp., *Escherichia vulneris*, *Mora staphylococcus*, *Moraxella atlantae*, *Moraxella fulton, Mucor* sp., *Proteus* sp., *Pseudomonas putida*, *Serratia marcescens*, *Streptococcus agalactiae*, *Staphylococcus caprae*, *Staphylococcus saprophyticus*, *Staphylococcus warneri*, *Viridans streptococci* and *Weeksella virosa*.

Different numbers of samples were sent for detection of bacteria using bacterial cultures. Correlations between the number of samples sent and the detection rate were examined through pairwise comparisons (Table 5). Five samples sent to bacterial culture achieved highest detection rate.

### Identifying Pathogenic Bacteria

According to the results, it is statistically significant between 2 samples and 4 samples, 2 samples and 5 samples, 3 samples and 5 samples. Five samples sent for detection has the greatest likelihood of identifying pathogenic bacterial (Table 6).

### Sinus Tract

Samples collected from 58 patients with sinus tracts were sent for bacterial culture (Table 7).

We compared the result of sinus tract bacterial culture with the pathogenic bacteria. Only less than half of samples (42.1%) from sinus tract were in consistent with the pathogenic bacteria.

### Two-Stage Debridement and One-Stage Debridement

Two-stage debridement treatment has higher union rate (87.0%) than the one-stage debridement (71.4%) and the result was statistically significant (p<0.05). In addition, one-stage debridement has more recurrence than two-stage debridement (p<0.05).

## Discussion

### Analysis of General Data

#### Gender Ratio

Among patients with chronic osteomyelitis, the male:female ratio was 4:1. This is in accordance with Kremers’s report [10], which argued that increasing road and industrial accidents contributed to the greater number of male patients, since males are more likely to engage in heavy physical labor or high risk activities.

#### Age Distribution

The current study showed chronic osteomyelitis was highest in the age group of 41 to 50 year-olds (29.0%), probably because road accidents frequently occur among those aged 20-50 years [11]. Traffic trauma is a high energy trauma that often leads to an open wound, a potential pathogenic factor for traumatic osteomyelitis. In addition, this age group has a high incidence of diabetes, which increases the frequency of diabetes-related osteomyelitis. Kremers [10] reports that American patients with diabetes-related osteomyelitis have increased from 2.3/100,000 in the late 1970s, to 10.5/100,000 in the 1990s, although currently its incidence appears to be stable at approximately 7.6/100,000. According to Cierny et al. [12], immunity of an organism is a significant factor in the occurrence and transformation of osteomyelitis. Different age groups have different resistance to infection, and older individuals may be more likely to have osteomyelitis because they have weaker immune systems.

### Analysis of Disease Data

#### Infection Site

The top three infection sites of chronic osteomyelitis - the femur (30.2%), tibiofibular (25.9%) and hip joint (15.3%) – account for 71.4% of all infections. This is consistent with other clinical reports [13,14], but differs from the clinical reports of south China. Wang et al. [8] reports that in southwest China, the top two infection sites are tibia (57.5%) and femur (26.8%). Jiang et al. [9] reported a similar result for southern China, finding the most frequent single infection site was the tibia (39.00%), followed by the femur (24.46%), and calcaneus (11.46%). One possible reason for this difference between north and south China could be the dense population found in southern China, leading to more frequent road and industrial accidents. As noted above, such accidents are more likely to create trauma with an open wound, and the thin soft tissue surrounding the tibia and lack of blood supply can increase the likelihood of wound infection, followed by chronic osteomyelitis. Thus, tibia osteomyelitis has the highest incidence in the north China.

#### Cierny-Mader Classification

The top three classifications in the current study are type IIA (23.1%), type IIIA (20.4%), and type IVA (18.0%), accounting for 61.5% of cases in total. In the domestic literature, the Cierny-Mader classification information given is often insufficient, with only anatomical type staging provided and no classification of the physiological type of the host [15]. In foreign literature using the Cierny-Mader classification, different literature focuses on different case groups, and the results also differ from each other. Hence, there is no uniform consensus on the statistical distribution of *Cierny-Mader* types [16-18].

Cierny et al. [12] discussed osteomyelitis classification in detail. They held the view that osteomyelitis classifications are not fixed for a particular case, but rather that osteomyelitis may transform from one classification to another in the same patient, as a process of natural progression or treatment. Thus, different types of osteomyelitis can be thought of as different pathological/physiological stages of disease [12]. The current study suggests that most chronic osteomyelitis falls into types II, III, and IV. For the physiological class, host A is more common than host B, and host C type is seen only rarely.

#### Bacterial Culture

The results of debridement bacterial cultures indicate that g+ bacteria (65.3%) was more common than g- bacteria (33.9%). The five most commonly observed bacteria were *Staphylococcus aureus* (35.51%), *S. epidermidis* (14.29%), *Pseudomonas aeruginosa* (9.8%), *Enterobacter cloacae* (5.31%), and *Escherichia coli* (4.08%); these accounted for 69.0% of the infections observed. The two g+ *Staphylococcus* bacteria made up nearly 50% of infections. The remaining three of the most five common bacteria were all g- bacteria. This is similar to most results found in clinical reports in China and abroad [8, 9, 19, 20, 21].

#### Detection Rate for Different Numbers of Samples

Identifying the pathogenic bacteria involved is an important step in treating chronic osteomyelitis. Based on characteristics of the pathogenic bacteria, effective antibiotics specific to the bacteria found should be used in clinical practice; this is a precondition to guaranteeing a curative effect. Sending more samples at once was an effective method to improve the detection rate of pathogenic bacteria. Lew et al. [22] suggested that five samples or more should be sent when treating chronic osteomyelitis, and that this could significantly improve the microbial detection rate. Our study similarly showed that the more sample sent for bacterial culture, the higher the detection rate. Sending only one sample had the lowest detection rate (66.7%), while five samples gave the highest rate (90.6%). Therefore, five or more samples should be sent from debridement to improve detection rates.

#### Identifying Pathogenic Bacteria

Typically, pathogenic bacteria and contamination bacteria are both observed in the results from bacterial cultures. If the same bacteria is detected twice or more in the culture results, it is presumed to be the pathogenic bacteria [23]. Using this principle, contaminant bacteria can be distinguished from pathogenic bacteria in the results. If all cultures give negative results, or if only one or differing results are observed among samples, then pathogenic bacteria cannot be determined. The current study indicates that the more samples that are sent for examination, the higher the probability of identifying the pathogenic bacteria. The probability of identifying the pathogenic bacteria was significantly improved (82.5%) if five or more samples were sent for bacterial culture; this is higher than the rates reported from Wang (71.5%) and Kremers et al. (75%).

#### Bacterial Culture of Sinus Tract Sample

In the current study, results from sinus tract bacterial culture were consistent with pathogenic bacteria in only 42.1% of cases. This might be caused by contamination with nonpathogenic bacteria during the sampling process, or other naturally-occurring random factors [8, 24]. Clinicians may also be collecting these samples in a non-standard manner. Regardless, the results from sinus tract bacterial culture can only be used as a reference for finding antibiotics to which bacteria are susceptible, not as the basis of the diagnosis and treatment decision.

### Clinical Treatments

In treating chronic osteomyelitis, the patient’s overall condition and local lesion must be considered to accurately assess the patient’s state before making careful decisions regarding treatment. The published literature includes a number of clinical treatments for chronic osteomyelitis [25-28], such as radical debridement, bone transport, acute limb shortening, two-staged reconstruction, local antibiotic therapy (antibiotic-impregnated beads, spacers, cements, intramedullary nails) and soft tissue grafting (free flap, myocutaneous flap, skin graft). Nonetheless, a common standard of diagnosis and treatment has not been established. Two basic consensuses are present regarding the treatment of chronic osteomyelitis. The first is that debridement is key to all treatments. The second is that guaranteeing a curative effect requires identifying the pathogenic bacteria involved and the use of appropriate antibiotics. The site of bone defect should be treated with local antibiotic-loaded PMMA spacers. External fixation or internal fixation can stabilize fracture ends.

Cierny [29] and Forsberg et al. [30] repeatedly emphasize radical debridement as the foundation for treating chronic osteomyelitis. A thorough scrape is needed at the dead space, necrotic tissue and sinus tract, until fresh blood is exuded, indicating healthy tissue, and plenty of irrigation must be done. Results of the current study suggest a good curative effect (87%) is achieved by repeated radical debridement, and implantation of appropriate antibiotic bone cement beads in the local lesion, with external or internal fixation. The reasons are as follow. 1) In the two-stage debridement method, debridement is completed in a more thorough manner. Within one week, debridement is performed twice. The second debridement removes all possible contaminated tissues based on the first debridement, thus reducing the possibility of recurrence [22, 31-32]. 2) Sending five samples for bacterial culture in the first debridement improves the detection rate of pathogenic bacterial. Further, an appropriate (i.e., one the bacteria was susceptible to) antibiotic release-carrier can be identified; it is more successful to use antibiotics based on the results of the bacterial culture and susceptibility testing. This also ensures rare bacteria are quickly recognized, and reduces the side effects of antibiotics since targeted antibiotics are used [33]. 3) An appropriate antibiotic release-carrier implanted after one week maintains the effective blood concentration of the antibiotic in local tissue longer. Previous research has shown that, after the blood antibiotic concentration reached its peak, it would gradually decline, so that after a few weeks, local antibiotic blood concentration was less than the effective blood concentration [34, 35]. Therefore, implanting a second, targeted antibiotic release carrier after one week could extend the duration of the local blood concentration, thus better inhibiting the growth of pathogenic bacteria. The method of repeating radical debridement and local antibiotic-loaded PMMA spacers ± external fixation gave good clinical results and can be recommended. One drawback of this approach is that it increases the number of operations required and prolongs hospitalization, thus increasing expenses.

The present study had several limitations. Since it was conducted in a single medical center in north China, it may not characterize chronic osteomyelitis in the whole area. Therefore, multicenter studies should be performed to obtain more detailed data. Second, this study lacked detailed statistical data on oral antibiotic therapy because most patients received oral antibiotics out of hospital.

## Conclusions

In summary, our study showed that chronic osteomyelitis is more common in males and in the age group from 41-50 years of age. Common infection sites are the femur, tibiofibular, and hip joint. Using Cierny-Mader classification, type IIA, IIIA and IVA infections were most common. More g+ than g- bacterial infections were observed, with *S. aureus* the most commonly observed pathogenic organism.

To improve pathogen detection rate, five or more samples should be sent for bacterial culture. Bacterial culture of the sinus tract performed poorly in identifying pathogenic bacteria, so choosing the appropriate antibiotic based on the result of sinus tract bacteria is not advised. Repeated radical debridement and identification of pathogenic bacteria are keys to successful treatment of chronic osteomyelitis. The current study suggests a treatment of two stage debridement + antibiotic-loaded PMMA spacers ± external fixation is the most effective.

## Acknowledgment

This research received no specific grant from any funding agency in the public, commercial, or not-for-profit sectors. There is no conflict of interests in the current research.

---

**Table S1.**
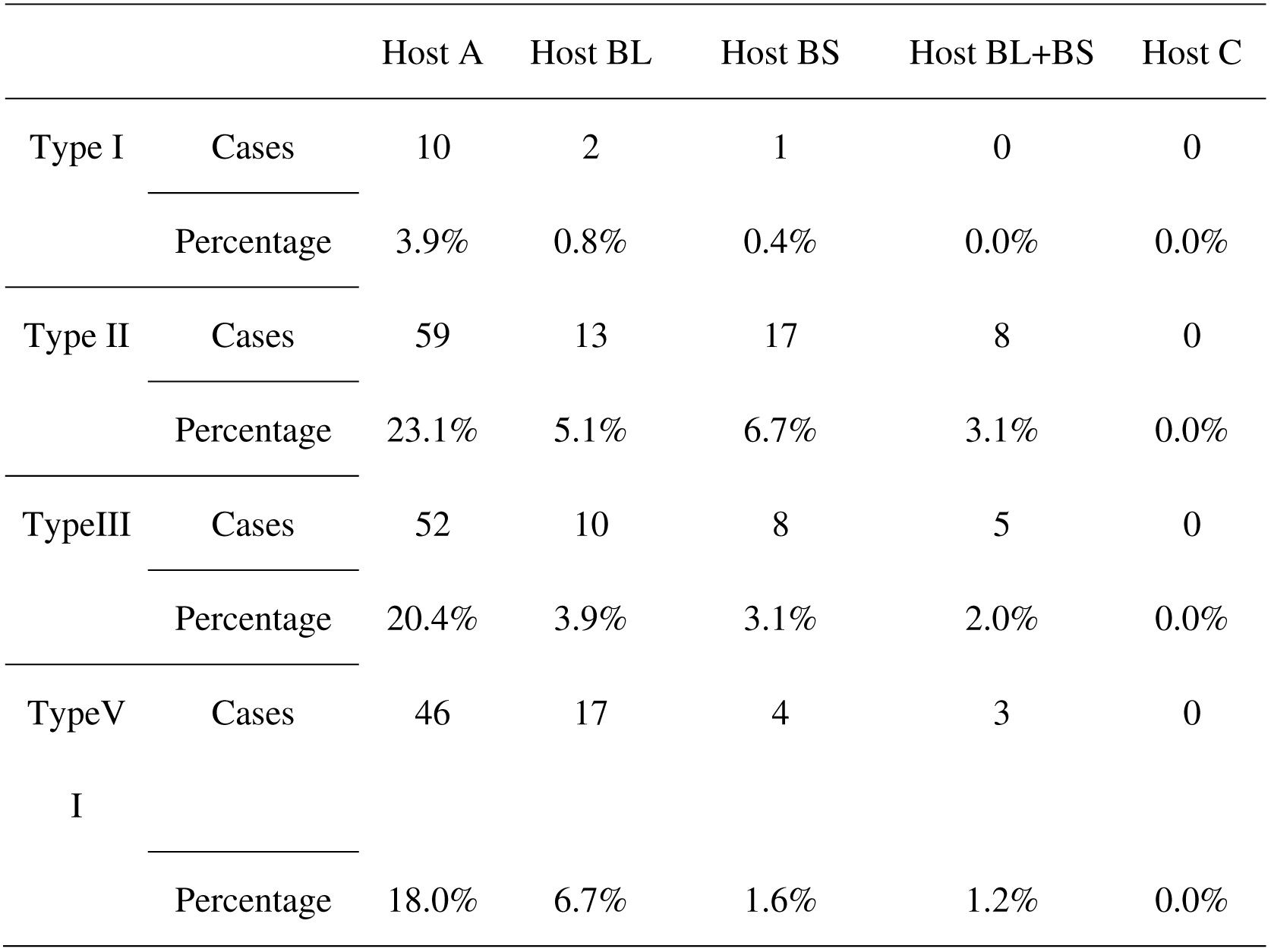
Cierny-Mader classification results.

**Table S2.** Laboratory tests of WBC, ESR and CRP.

|  | Frequency | Percentage |
| --- | --- | --- |
| WBC(+) | 104 | 32.2% |
| WBC(-) | 219 | 67.8% |
| ESR(+) | 230 | 71.2% |
| ESR(-) | 93 | 28.8% |
| CRP(+) | 290 | 89.7% |
| CRP(-) | 83 | 10.3% |

**Table S3.** Comprehensive statistics of WBC, ESR and CRP.

|  | WBC | ESR | CRP | Times | Percentage |
| --- | --- | --- | --- | --- | --- |
| Result 1 | (-) | (-) | (-) | 20 | 6.6% |
| Result 2 | (+) | (-) | (-) | 3 | 1.0% |
| Result 3 | (-) | (+) | (-) | 2 | 0.7% |
| Result 4 | (-) | (-) | (+) | 53 | 17.4% |
| Result 5 | (+) | (+) | (-) | 0 | 0.0% |
| Result 6 | (+) | (-) | (+) | 8 | 2.6% |
| Result 7 | (-) | (+) | (+) | 132 | 43.4% |
| Result 8 | (+) | (+) | (+) | 86 | 28.3% |
| In total |  |  |  | 304 | 100.0% |

**Table S4.** Bacterial distribution.

| Bacterial | Frequency | Percentage |
| --- | --- | --- |
| Staphylococcus aureus | 87 | 35.51% |
| Staphylococcus epidermidis | 35 | 14.29% |
| Pseudomonas aeruginosa | 24 | 9.80% |
| Enterobacter cloacae | 13 | 5.31% |
| Escherichia coli | 11 | 4.49% |
| Klebsiella pneumoniae | 7 | 2.86% |
| Staphylococcus capitis | 6 | 2.45% |
| Baumann's acinetobacter complex. | 6 | 2.45% |
| Enterococcus faecalis | 5 | 2.04% |
| Staphylococcus haemolyticus | 4 | 1.63% |
| Enterococcus faecium | 4 | 1.63% |
| Staphylococcus hominis | 4 | 1.63% |
| Serratia marcescens | 4 | 1.63% |
| Others |  | 14.28% |

**Table S5.** Detection rate when using different numbers of samples, and the correlations between detection rate and number of samples.

| Number of samples for detection | Detection times | Negative result | Positive result | Detection rate | Compare with other groups | P Value |
| --- | --- | --- | --- | --- | --- | --- |
| 1 sample | 63 | 21 | 42 | 66.7% | VS 2 sample | >0.05 |
|  |  |  |  |  | VS 3 samples | >0.05 |
|  |  |  |  |  | VS 4 samples | >0.05 |
|  |  |  |  |  | VS 5 samples | <0.05 |
| 2 samples | 76 | 23 | 53 | 69.7% | VS 3 samples | >0.05 |
|  |  |  |  |  | VS 4 samples | >0.05 |
|  |  |  |  |  | VS 5 samples | <0.05 |
| 3 samples | 51 | 13 | 38 | 74.5% | VS 4 samples | >0.05 |
|  |  |  |  |  | VS 5 samples | <0.05 |
| 4 samples | 41 | 9 | 32 | 78.0% | VS 5 samples | >0.05 |
| 5 samples or above | 64 | 6 | 58 | 90.6% |  |  |

**Table S6.**
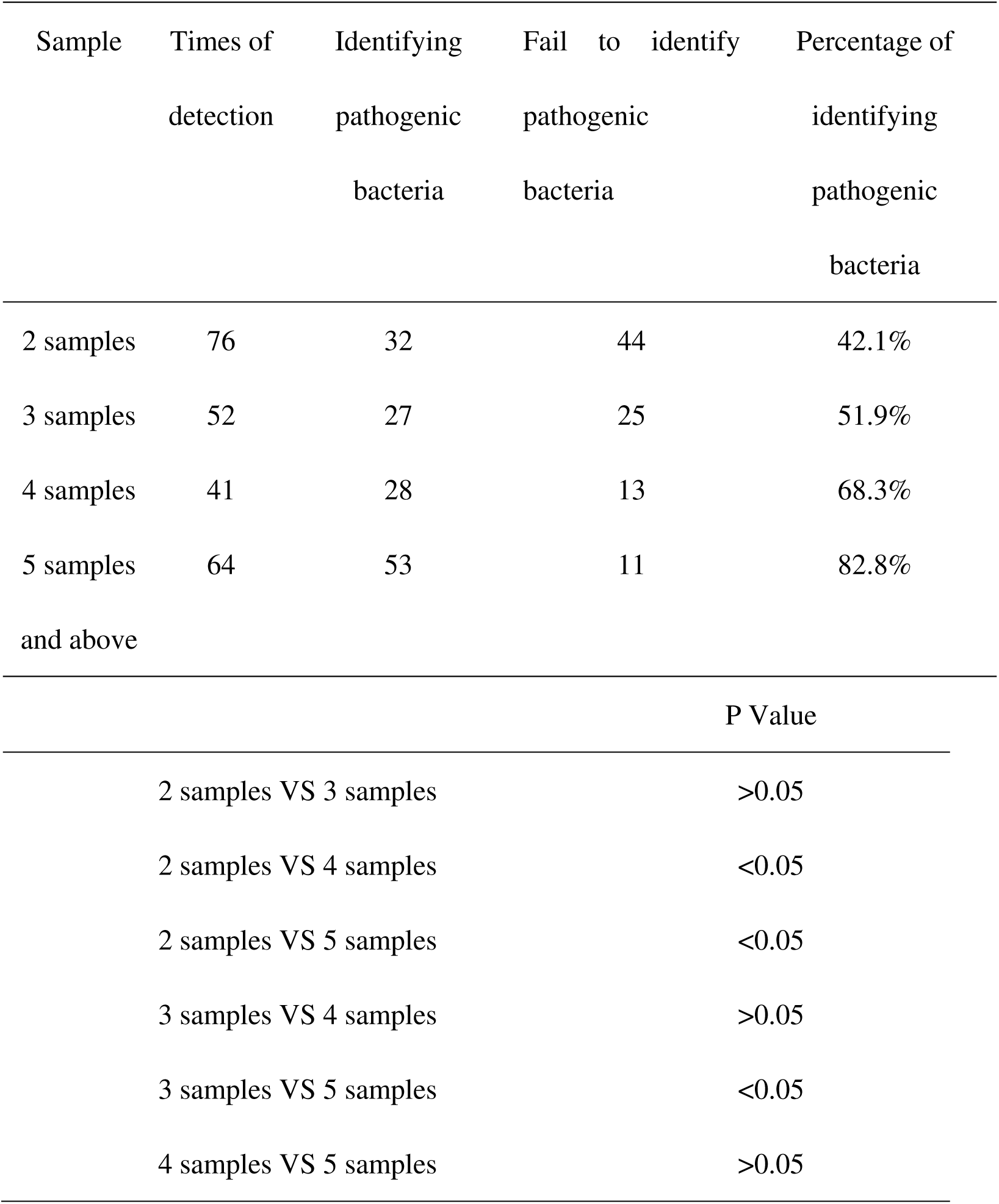
Ability to identify pathogenic bacteria for different numbers of sample, and the correlation between number of samples and identification success.

**Table S7.** The distribution of bacteria found in the sinus tract.

| Bacteria | Frequency | percentage |
| --- | --- | --- |
| staphylococcus aureus | 15 | 25.86% |
| staphylococcus epidermidis | 9 | 15.52% |
| pseudomonas aeruginosa | 5 | 8.62% |
| enterobacter cloacae | 3 | 5.17% |
| escherichia coli | 3 | 5.17% |
| staphylococcus haemolyticus | 2 | 3.45% |
| enterococcus faecalis | 2 | 3.45% |
| klebsiella pneumoniae | 2 | 3.45% |
| Baumann's acinetobacter<br>complex. | 2 | 3.45% |
| Candida albicans | 2 | 3.45% |
| Rhodococcus rhodococcus | 2 | 3.45% |
| peptostreptococcus anaerobius | 2 | 3.45% |
| staphylococcus warneri | 1 | 1.72% |
| Enterobacter agglomerans | 1 | 1.72% |
| enterobacter | 1 | 1.72% |
| providencia rettgeri | 1 | 1.72% |
| staphylococcus hominis | 1 | 1.72% |
| citrobacter braakii | 1 | 1.72% |
| staphylococcus saprophyticus | 1 | 1.72% |
| proteus mirabilis | 1 | 1.72% |
| klebsiella oxytoca | 1 | 1.72% |
| In total | 58 | 100.0% |

**Table S8.** The coincidence rate between the result from sinus tract bacterial culture and pathogenic bacteria.

|  | Numbers | Percentage |
| --- | --- | --- |
| Total Positive samples | 57 | 100.0% |
| Same with pathogenic bacteria | 24 | 42.1% |
| Different with pathogenic bacteria | 33 | 57.9% |

**Table S9.** Clinical effects and prognosis of the two different treatments.

| Treatment | Two-stage<br>(108cases ) | one-stage<br>(147cases ) | P-value |
| --- | --- | --- | --- |
| union | 94 | 105 | <0.05 |
| Recurrence | 14 | 42 | <0.05 |
| Union rate after first treatment | 87.0% | 71.4% | <0.05 |
| Continue the treatment after<br>Recurrence | 11 | 30 | >0.05 |
| Union after once retreatment | 9 | 4 | <0.05 |
| Union after twice retreatments | 0 | 13 | - |
| Union after thrice treatments | 0 | 12 | - |
| Withdraw treatment after<br>Recurrence | 3 | 6 | >0.05 |
| Nonunion(amputation) | 2 | 1 | >0.05 |

